# The Asian gypsy moth (*Lymantria dispar* L.) populations: resistance of eggs to extreme winter temperatures

**DOI:** 10.1101/2021.02.09.430420

**Authors:** G.G. Ananko, A.V. Kolosov

## Abstract

Gypsy moth *Lymantria dispar* (GM) is a polyphagous insect and one of the most significant pests in the forests of Eurasia and North America. Accurate information on GM cold hardiness is needed to improve methods for the prediction of population outbreaks, as well as for forecasting possible GM range displacements due to climate change.

As a result of laboratory and field studies, we found that the lower lethal temperature (at which all *L. dispar asiatica* eggs die) range from –29.0 °C to –29.9 °C for three studied populations, and no egg survived cooling to –29.9 °C. These limits agree to within one degree with the previously established cold hardiness limits of the European subspecies *L. dispar dispar*, which is also found in North America. This coincidence indicates that the lower lethal temperature of *L. dispar* is conservative.

Thus, we found that the Siberian populations of GM inhabit an area where winter temperatures go beyond the limits of egg physiological tolerance, because temperature often fall below –30 °C. Apparently, it is due to the flexibility of ovipositional behavior that *L. dispar asiatica* survives in Siberia: the lack of physiological tolerance of eggs is compensated by choosing warm biotopes for oviposition. One of the most important factors contributing to the survival of GM eggs in Siberia is the stability of snow cover.

**Summary:** Within the geographical range of Siberian gypsy moth populations, extreme temperatures go beyond the limits of the physiological tolerance of wintering eggs (–29.9 °C), and their survival depends on the choice of warm biotopes for oviposition.

## INTRODUCTION

Gypsy moth *Lymantria dispar* L. 1758 *(Lepidoptera: Erebidae)* is a polyphagous insect and one of the most important pests in forestry, because periodical outbreaks can cover hundreds of thousands of hectares of forest areas (Qian, 2000; Ponomarev et al., 2012). The role of various factors determining the dynamics of gypsy moth (GM) population outbreaks remains debatable (Liebhold et al., 2000; Ponomarev et al., 2012; Martemyanov et al., 2019). Another issue is related to the observed climate changes (IPCC, 2013), which are particularly pronounced in the circumpolar regions of Siberia: how will the Siberian GM populations respond to warming? will there be a shift of the northern boundary of their range? Climate warming can change species distribution and population cycles, but the mechanisms of these processes are yet unclear (Klapwijk et al., 2012; Myers and Cory, 2013). Evidence suggests that climate warming can lead to significant changes in population outbreak dynamics and amplitude (Esper et al. 2007; Jepsen et al. 2008).

Predicting gypsy moth outbreak dynamics and possible shifts of the northern border of its range is impossible without accurate information about the insect’s physiological tolerance limits. GM overwinter as eggs deposited in a variety of locations, so it is particularly interesting to study the cold hardiness of the eggs. There is a significant body of research on egg wintering conditions of European and North American populations of *L. dispar dispar* (Pogue and Schaefer, 2007; Sullivan and Wallace, 1972; Leonard, 1974; Madrid and Stewart, 1981), and new studies appear periodically (Streifel et al., 2019; Fält-Nardmann et al., 2018). According to different sources, the supercooling point of *L. dispar dispar* eggs varies in the range from –23.6 °C to –30.3 °C. However, the cold hardiness of eggs of Asian populations of the gypsy moth *L. dispar asiatica* Vnukovskij (Pogue and Schaefer, 2007, revised status) has not been sufficiently studied. Within the subspecies *L. dispar asiatica* there are four geographical forms that differ mainly in the ovipositional behavior of females (Ponomarev et al, 2012). In Russia, the West Siberian, East Siberian and Far Eastern geographical forms are found, and the Central Asian GM form is found in the countries of Central Asia, particularly in Kyrgyzstan. There is evidence that the eggs of Asian populations are more resistant to low temperatures than those of European populations. For example, in studying the cold hardiness of the North Caucasian and Far Eastern GM populations, it was found that eggs die at a lower temperature, about –45… –48 °C (Pantyukhov, 1964). According to long-term meteorological observations (www.pogodaiklimat.ru) in the habitat of *L. dispar asiatica* (in Siberia), the temperature often drops to –30… –42 °C during a typical winter, but values below –48 °C were recorded only a few times during the entire observation period (since 1959). At the same time, Russian researchers of Siberian (Il′ynskiy and Tropin, 1965; Ponomarev et al, 2012) and Far Eastern (Yurchenko and Turova, 1984) GM populations often note significant egg mortality in winter. Thus, the eggs of Siberian populations often die, despite that the temperature does not fall to –48 °C. It remains unclear whether the eggs die of a short-term decrease of winter temperature below the physiological tolerance limit or they (also) die as a result of prolonged exposure at sublethal temperatures? and do the eggs of different Asian populations differ in cold hardiness?

To answer these questions, we investigated the resistance of GM eggs to extremely low temperatures in laboratory and field conditions. The main objective of the study is to determine the lower temperature hardiness limit of the Novosibirsk, Altai, and Kyrgyz populations belonging to the West Siberian, East Siberian, and Central Asian geographical forms of *L. dispar asiatica*, respectively.

## MATERIALS AND METHODS

### Samples

We studied eggs of three Asian populations of *Lymantria dispar*. These populations belong to three geographical forms showing significant ecological differences (Ponomarev et al., 2012).

<u>The Novosibirsk population</u> belongs to the West Siberian geographical form. Egg masses were sampled in September 2019 in the Ordynsky district of the Novosibirsk region, Russia (54.22° N, 81.89° E). The range of the West Siberian geographical form spans forests from Ural to Altai and Sayan mountains. The northern range of the West Siberian geographical form is unstable and is moving northbound across the deciduous forests of the West Siberian plain due to the natural climate changes occurred in the 20^th^ century. Females oviposits at the host trees: the mean height at which eggs are placed is 6-7 cm above the ground level (Ponomarev et al., 2012).

<u>The Altai population</u> belongs to the East Siberian geographical form. Egg masses were collected in September 2019 in the Ust’-Kansky district of the Altai Republic, Russia (51.33° N, 84.74° E). The range of the East Siberian geographical form is vast and spans deciduous and larch forests from Altai in the west to the southern foothills of Stanovoi Range in the east, as well as Mongolia and Khingan. One of the characteristic features of this geographical form is that females usually do not oviposit on tree trunks and rather place them on rock outcrops (Benkevich, 1956).

<u>The Kyrgyz population</u> belongs to the Central Asian geographical form. Egg masses were collected in the Nookensky district of the Jalalabad region of the Kyrgyz Republic (41.10° N, E 72.58° E). Its extensive range includes southern areas of Kazakhstan, as well as Kyrgyzstan, Uzbekistan, Tajikistan, and also apparently Turkmenistan, Afghanistan and Xinjiang-Uyghur Autonomous Region of China. In forests of South Kyrgyzstan, females oviposit in tree hollows and bark cracks, on stones and ground (Romanenko, 1981). Egg masses on tree trunks can be found at a height from the ground level to several meters above the ground (the authors’ own observations).

### Determination of egg viability

To stimulate larvae hatching, samples of eggs cleaned of the hair coating (100 pcs.) were placed into a 10-cm Petri dish. Eggs were incubated under the following conditions: 26 °C, relative humidity 60 %, day/night regime (16/8 h). We counted the hatched larvae every day and continued the experiment for one week after all larvae hatched, but not less than 4 weeks. Experiments were repeated in 3 replicates with 100 eggs in each. Viability of eggs from original (i.e., unexposed to negative temperatures) samples was different depending on population: 96 % for the Novosibirsk, 82 % for the Altai, and 78 % for the Kyrgyz population. In order to facilitate the comparison of cold hardiness of eggs in different gypsy moth populations, we normalized all the obtained values as if the viability of original eggs (before the experiment) were 100 % in all cases.

### Determination of egg cold hardiness in laboratory conditions

Egg masses were stored at 6 °C. Immediately prior to experiments, eggs were cleaned of the hair coating and packed in paper bags, 100 eggs per each. To let eggs adapt to the cold, the heat insulating chamber containing the samples was placed to freezer which was set at –15 °C. On the day of experiment, the sample chamber was transferred to a low-temperature freezer set at –35 °C. During the experiment, eggs were gradually (1–2 degrees per hour) cooled to a desired temperature. Cooling of samples from –15 °C to target temperature took 11–14 hours. Only one temperature value per experiment was tested, because once the preset target temperature was reached, the chamber was immediately removed from the freezer and left at room temperature in order to ensure a slow heating of the samples.

Three egg samples belonging to a certain gypsy moth population, 100 pcs. each, were used per each experimental point. We placed three identical samples into a plastic test tube, so that they were in direct contact with a temperature sensor connected by a cable to an external temperature recorder. The test tube with samples and sensor was placed into a special chamber for the slow cooling of samples, which consisted of heat insulating chamber with hollow walls filled with 60 % glycerin solution; we added an additional heat insulation by foam plastic and cottonwool inside the chamber. The sample chamber was cooled in a low-temperature freezer, and sample temperature was sent along the wire to the logger, which was placed next to the freezer, every 30 seconds. Temperature was recorded using the 2-channel EClerk-M-11-2Pt-HP logger (Relsib, Russia: https://relsib.com/product/izmeritel-registrator-temperatury-eclerk-m-2pt-hp) with measurement accuracy of 0.2 °C.

### Field technique for the assessment of egg cold hardiness

Short 1–5-day field studies were carried out in winter 2019/2020. When a significant cooling was forecast, we attached egg samples to a birch trunk at a certain height: 10 cm above the ground (under a snow layer), and 100 cm above the ground (i.e., above the snow surface). After the end of the cold weather, eggs were tested for viability. Three egg samples belonging to a certain gypsy moth population, 100 pcs. each, were used per each experimental point. Each sample was wrapped in filter paper. Samples were placed into a cotton bag to prevent egg eating by birds. Temperature sensor was put into a similar paper bag next to egg samples. Sample temperatures were recorded every 10 minutes.

### Statistical evaluation of the results

We used the STATISTICA Advanced software package (http://statsoft.ru/products/STATISTICA_Advanced/). Graphs and tables show average values, calculated from three replicates per point. Data shown as mean ± standard error of the mean (mean ± s.e.m.).

## RESULTS

### Preliminary studies

Preliminary field studies of gypsy moth egg cold hardiness were conducted during the winter season from November 2018 till March 2019. For the experiments carried out in the neighbourhood of Novosibirsk, we used egg masses of the local Novosibirsk population, which belongs to the West Siberian geographical form. We attached samples to a birch trunk at a certain height. The first batch of samples was put 10 cm above the ground level, under a snow layer (snow depth 0–70 cm), and the second batch was placed 100 cm above the ground level (i.e., above the snow surface). Samples were tested for viability twice: in January and in March. We have found that in the egg masses that overwintered under the snow, the viability rate remained almost the same, in contrast to the samples overwintered in open air, where all eggs died already by the moment of the first sampling in mid-January. According to the weather data obtained by the local station [www.pogodaiklimat.ru], which is 12 km from the experimental site, the temperature fell below –30 °C five times during the period from November 2018 to the 15^th^ of January, 2019, with a minimum of –34,9 °C. Winter temperatures in this period were typical for the region; nevertheless, eggs of the local GM population died already by the middle of the winter.

In the next winter season (2019/2020) we carried out a series of laboratory and field studies of cold hardiness of as many as three Asian gypsy moth populations: the Novosibirsk, Altai, and Kyrgyz populations. Continuous temperature recording on experimental sites was organized to ensure accurate results.

### Laboratory testing of GM egg cold hardiness

In order to assess cold hardiness of gypsy moth eggs, we carried out a series of experiments with sample cooling in a low-temperature freezer. Samples were put in a special chamber and cooled to a desired temperature (at a rate of 1–2 degrees per hour), whereafter the chamber was immediately removed from the freezer. We examined the temperature range from –23 °C to –29 °C with 1 °C step, and the temperature range from –29 °C to –29.9 °C with 0.3 °C step. The obtained data (Fig. 1) indicate obvious heterogeneity of the egg sample in terms of cold hardiness: a minor part of the eggs dies already at –24 °C, and no eggs survive if the temperature drops to –29,9 °C. A possible cause of the observed differences in egg cold resistance may be that a mixture of eggs from many family egg masses was used in the experiments. Use of a random egg sample from many different egg masses provides more accurate estimates of a population’s cold hardiness limits, compared to experiments with individual egg masses.

**Figure 1.**
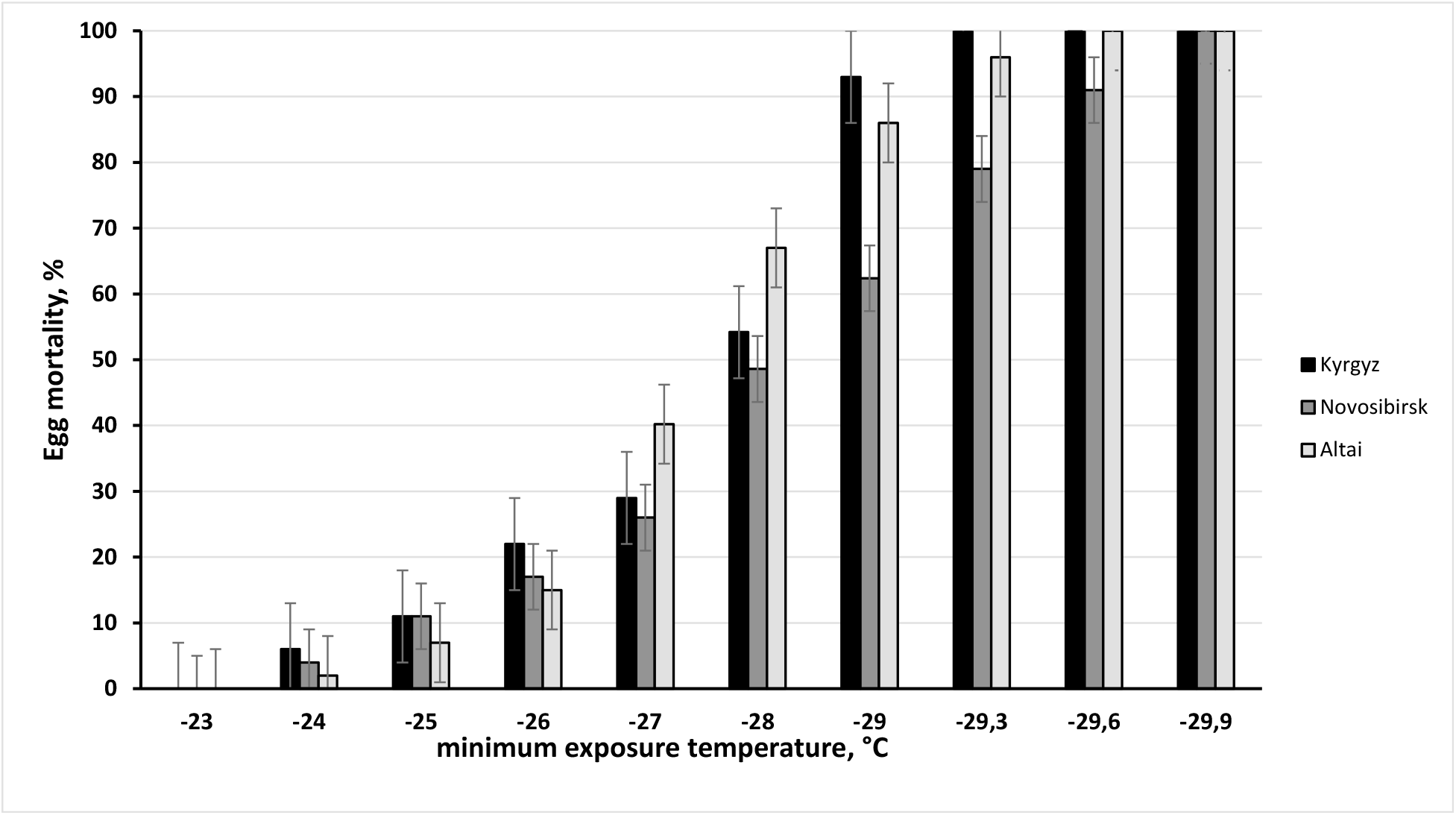
Mortality dynamics (mean ± s.e.m.) of gypsy moth eggs under cooling to extreme temperatures.

The temperature at which the death of 50 % of eggs was observed, was –(27.8±0.2) °C, – (28.l±0.2) °C, and –(27.4±0.2) °C for the Kyrgyz, Novosibirsk, and Altai populations, respectively. We repeated the experiment with cooling of samples belonging to the three populations to –29.9 °C a number of times, but the result was always the same: all eggs died. It is noteworthy that the temperature was between –29.6 and –29.9 °C for as little as 14 minutes. Hence, even a short-term temperature drop to the critical value of –29,9 °C leads to the death of 100 % of eggs. It is also noteworthy that the temperatures at which 100 % of eggs die are not substantially different for the three geographically distant populations, ranging from –29,3 °C in the Kyrgyz population to –29.9 °C in the Novosibirsk population. The lower cold hardiness limit of the studied Asian populations is apparently a conservative value, since the ascertained inter-population difference is less than one degree.

### Field studies of GM egg cold hardiness

In order to validate the results of laboratory tests, we designed short-term (1–5-day) field experiments. Based on weather forecast, before the nearest cold snap we arranged egg samples on a birch tree trunk, and after the cold snap the samples were tested for egg viability.

In the field experiment No. 1, which lasted around 5 days (December 25–30, 2019), we carried out a comparative study of two biotopes (see the temperature plot on the Fig. 2). The first sample batch was attached to a birch trunk at the height of 10 cm above the ground, so that a 70-cm-high snow layer covered the samples. The second sample batch was fixed on the same birch trunk at the height of 100 cm above the ground (i.e., above the snow surface).

**Figure 2.**
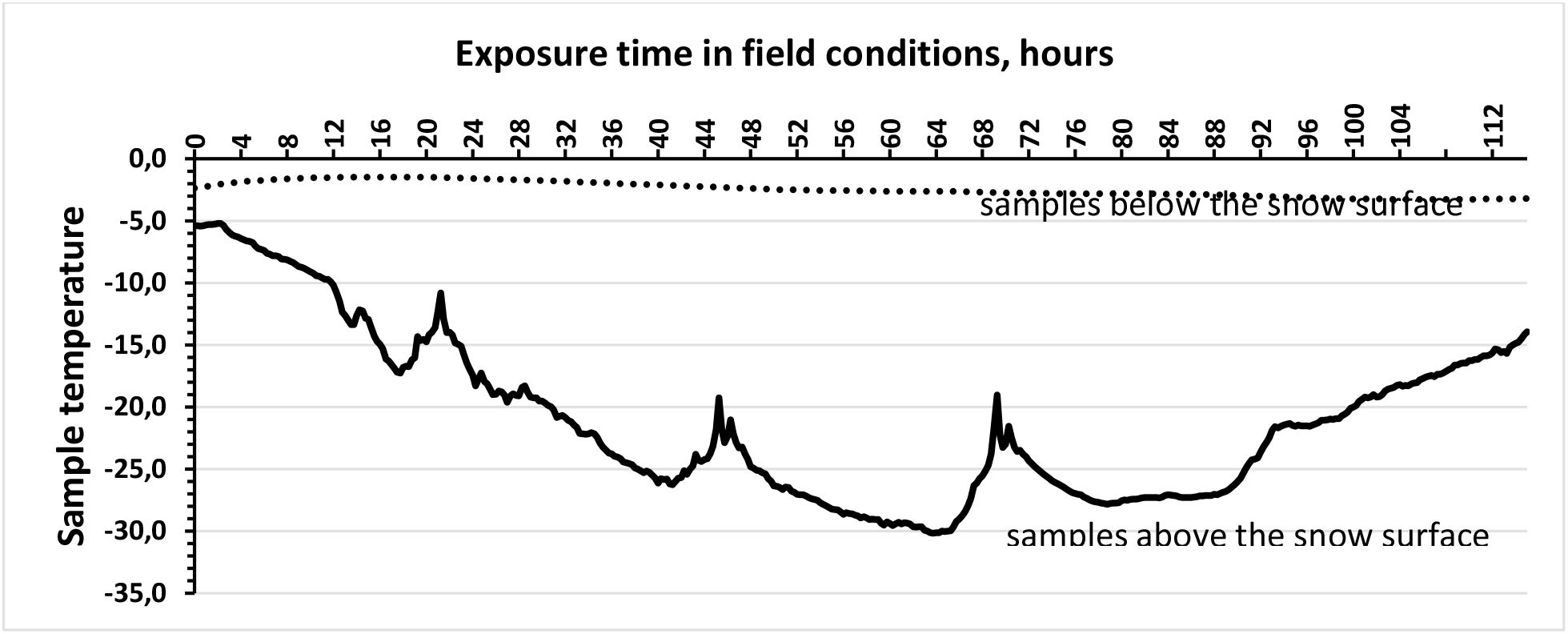
Temperature as a function of time during the field experiment No. 1. Samples of eggs of the Novosibirsk, Altai, and Kyrgyz populations were exposed on a birch trunk in 2 locations, the first at the height of 10 cm above the ground, so that a 70-cm-high snow layer covered the samples; the second above the snow surface, at the height of 100 cm above the ground.

Table 1 shows that the fate of eggs depends critically on the presence of snow cover. The viability of eggs covered with snow remained practically the same during the experiment. At the same time, all eggs died in samples fixed just 90 cm higher, but not covered with snow. It should be noted that the temperature fell below –30 °C for as little as 2 hours (Fig. 2). The results of this field research clearly confirm the laboratory tests: even a short-term temperature drop below the critical threshold of –30 °C kills the eggs of all the three populations in question.

**Table 1.** Results of the field experiment No. 1: egg mortality after the exposure under extreme below-freezing temperatures.

| Population | Egg mortality (mean ± s.e.m.), %, after exposure in field conditions at minimum temperature: | |
| --- | --- | --- |
| | $T_{\min} = -30.2\text{ }^{\circ}\text{C}$ | $T_{\min} = -3.3\text{ }^{\circ}\text{C}$ (under the snow) |
| Novosibirsk | $100 \pm 0$ | $2 \pm 1$ |
| Altai | $100 \pm 0$ | $1 \pm 1$ |
| Kyrgyz | $100 \pm 0$ | $3 \pm 2$ |

During the field study No. 2 (January 29–31, 2020), sampling was performed twice: after 20 and 42 hours of exposure in field conditions. The first sample batch was subjected to minimum temperature of –(26.1±0.2) °C (Fig. 3). The second sample batch, which was exposed for 42 hours, was subjected to extreme cooling twice: during the first day, there was a cold snap to –(26.1±0.2) °C, and an additional cooling down to –(28.7±0.2) °C after 20 hours.

**Figure 3.**
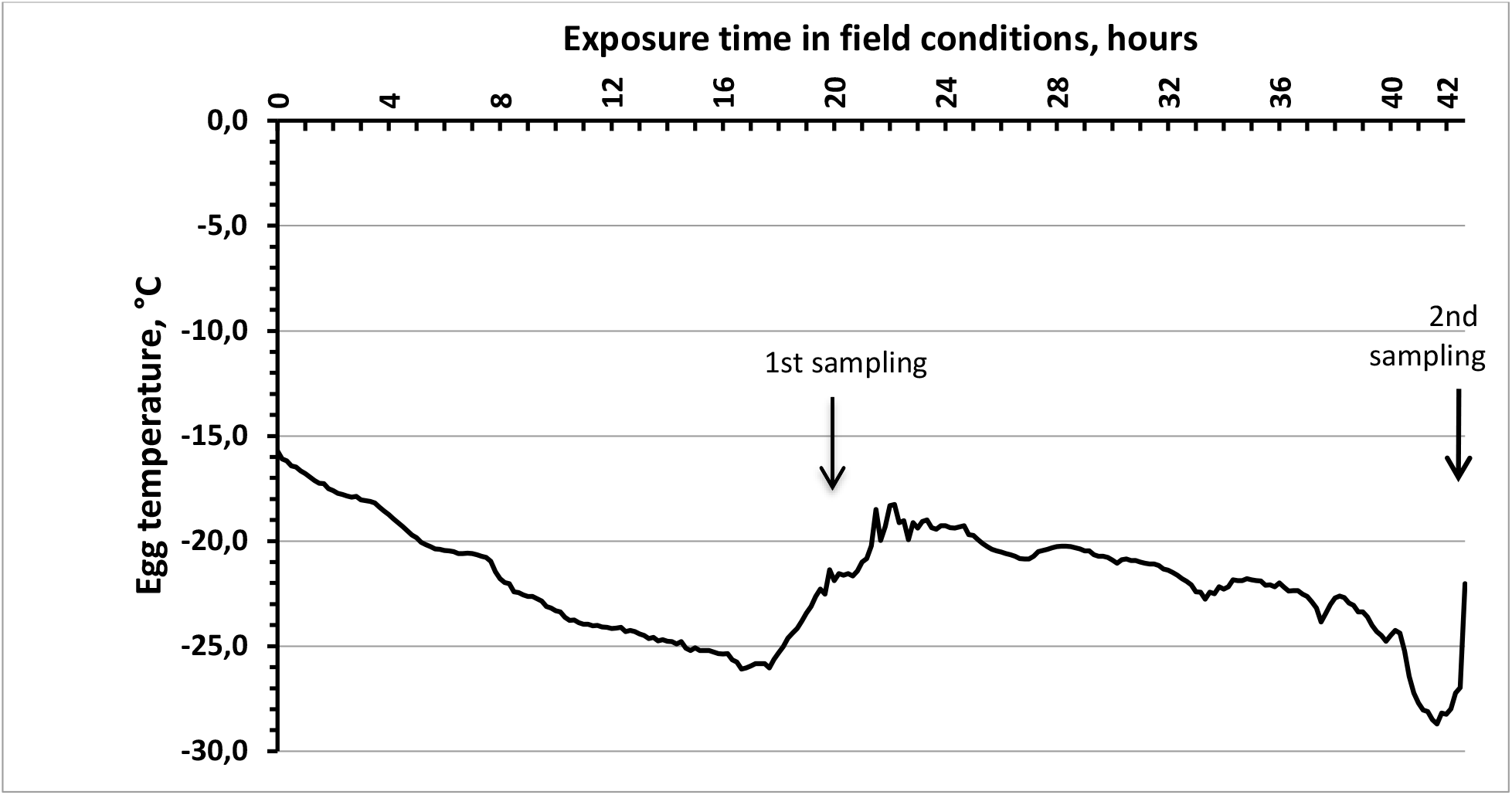
Temperature as a function of time during the field experiment No. 2. Samples of eggs of the Novosibirsk, Altai, and Kyrgyz populations were exposed above the snow surface. Eggs were sampled twice: after the first (20 hours of exposure) and second (42 hours of exposure) cold snap.

In general, the results of field studies (Table 2) corroborate the conclusions drawn from the laboratory experiments. However, the details of the conditions of laboratory and field experiments are, of course, different, such as in terms of exposure duration and temperature. We can see that in field conditions, under the minimum exposure temperature of –26.1 °C, more eggs died than in low-temperature freezer at –26.0 °C. For example, in the Kyrgyz population, 22 % of eggs died in the laboratory experiment, while in the field study the mortality was as high as 52 %. The other two populations also show similar differences. The mortality differences can probably be explained by different sample exposure durations: in the laboratory experiment, the impact of the minimum temperature was transient (ca. 5 minutes), whereas in field conditions, samples were exposed to the temperature of –26 °C for approximately an hour.

**Table 2.** Results of the field experiment No. 2: egg mortality after the exposure at extreme negative temperatures.

| Population | *Egg mortality (mean ± s.e.m.), %, after exposure in field conditions at minimum temperature: | |
| --- | --- | --- |
| | $T_{\min} = -26.1\text{ }^{\circ}\text{C}$ | $T_{\min} = -28.7\text{ }^{\circ}\text{C}$ |
| Novosibirsk | $39 \pm 4$ | $72 \pm 5$ |
| Altai | $22 \pm 3$ | $88 \pm 6$ |
| Kyrgyz | $52 \pm 5$ | $95 \pm 4$ |

\* Eggs were sampled twice: after 20 and 42 hours of exposure (temperature graph see in the Fig. 2).

### Prolonged exposure of GM eggs to the sublethal temperature

We have also studied the time dependency of egg viability during a prolonged exposure to the sublethal temperature, – 20 °C. Samples of the eggs of the Kyrgyz, Novosibirsk, and Altai populations were placed into a freezer, where the temperature of –(20 ± 2) °C was maintained. In order to insure gradual (at a rate of 1–2 degrees per hour) sample cooling from 6 °C дo –20 °C, we used a heat insulating chamber. We continued the experiment for 2 months until it was clear that all the eggs died. Samples were taken every 5 days; three samples, 100 eggs in each, were taken per experimental point and tested for viability. The test results have shown that there is progressive death of GM eggs during the prolonged exposure to the temperature of –20 °C (Fig. 4).

**Figure 4.**
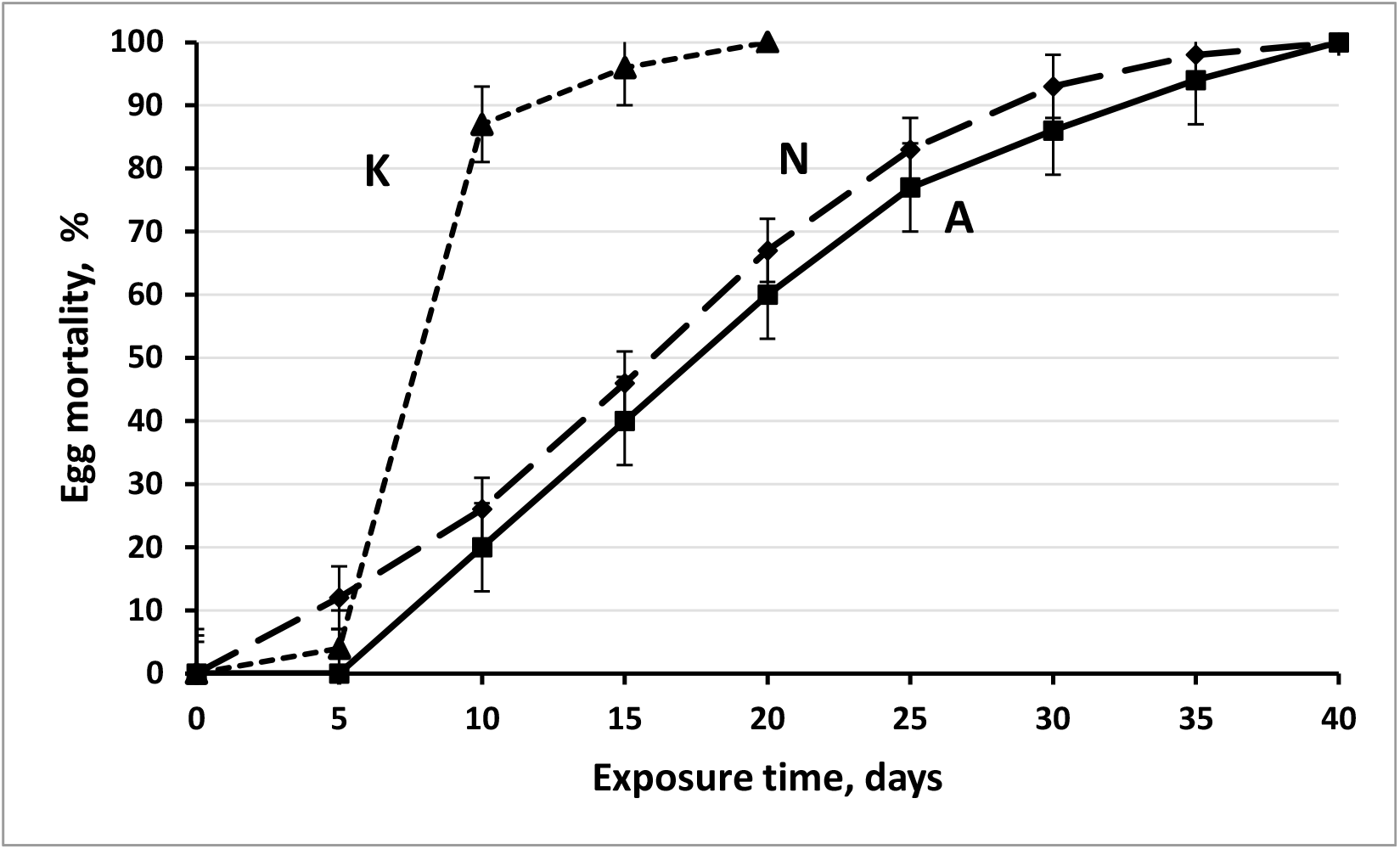
Mortality dynamics (mean ± s.e.m.) for the eggs of the Kyrgyz (K), Novosibirsk (N), and Altai (A) populations during prolonged exposure to the sublethal temperature, –(20 ± 2) °C.

The mortality dynamics is approximately the same for the eggs of the Novosibirsk and Altai populations, 100 % of eggs dying after a 40-day exposure. Eggs of the Kyrgyz population die much earlier: 90 % were dead as early as after the 10^th^ day, and there were no viable eggs after 20 days. Half of the eggs of the Kyrgyz, Novosibirsk, and Altai populations die after 7.8, 16.0, and 17.5 days of exposure at –20°C, respectively. Hence, eggs of the Central Asian geographical from (the Kyrgyz population) are significantly less resistant to prolonged exposure to sublethal temperatures than those of the West Siberian (the Novosibirsk population) and the East Siberian (the Altai population) geographical forms.

## DISCUSSION

During the preliminary studies carried out in winter 2018/2019 it was found that the eggs of the Novosibirsk GM population die as early as after 2 months of exposure to winter temperatures typical for Siberia. On the other hand, the viability of samples overwintered under the snow was not affected to any significant extent. During the next winter season (2019/2020) we have expanded the studies to include three populations: the Novosibirsk, Altai, and Kyrgyz populations, which belong to the West Siberian, East Siberian, and Central Asian geographical forms, respectively. Laboratory and field cold hardiness tests were carried out in parallel for these populations, and was temperature recorded exactly at locations where samples were left.

In modeling natural conditions, it is very important to choose adequate sample cooling rates (Terblanche et al., 2011). In doing that, we based on daily temperature fluctuation data and went on assumption that the GM egg masses are accustomed to typical daily temperature fluctuations during the Siberian winter. According to our observations, the typical rate of air temperature change is between 0 and about 2 degrees per hour, but in the morning and in the evening this rate may be significantly higher. We have chosen the slow sample cooling rate; heat insulation in the chamber was so designed that the sample cooling rate was not greater than 2 degrees per hour when the chamber was placed into the low-temperature freezer.

The resistance of insect eggs to below-freezing temperatures can be variable during winter and drops before larvae hatching (Madrid and Stewart, 1981). In Siberia, steady below-freezing winter temperatures are observed since November. Hence, it is by this date that GM eggs should attain the maximum cold hardiness. This is why the above reported series of experiments was conducted between November 2019 and January 2020.

Gypsy moth eggs, just as eggs of many other temperate-zone insect species, use the strategy of cold hardiness known as freeze avoidance. Stability of insect eggs in the supercooled state is favoured by small size, the absence of nucleation sites (Bale 2002, 2010), as well as by the presence of a cryoprotectant, such as glycerin, in the GM eggs (Denlinger et al., 1992).

According to different sources, supercooling points (SCP) for the eggs of the European form of *L. dispar* dispar, also found in North America, lie in the interval from –23.6 °C to – 30.3 °C (Sullivan and Wallace, 1972; Leonard, 1974; Madrid and Stewart, 1981; Fält-Nardmann et al., 2018). For instance, Waggoner (1985) found that a significant part of eggs freezes at temperatures below –26 °C. In a recent study (Fält-Nardmann et al., 2018) it was found that the median supercooling points of the German GM population vary broadly: from –23.6 to –24.4 °C in one egg batch, and from –27.7 to –28.9 °C in another. It should be noted that the mentioned range of the median supercooling points in the German GM population nearly coincides with that obtained in our study: eggs of Asian populations died in the broad temperature range from – 24.0 to –29.9 °C. Fält-Nardmann et al. argue that the northern range of the European GM form might reflect to some extent the physiological tolerance limit of wintering eggs. According to the data presented in that publication, the following correlation is observed: the northern range of the European GM populations approximately corresponds to the latitude where extreme winter temperatures do not fall below – 30 °C. Thus, the northern range roughly corresponds to the egg cold hardiness of the European GM populations. On this basis the authors forecast a possible northbound expansion of the GM range promoted by the increase of winter temperatures. Correlations between range and physiological cold hardiness limits are reported for other insect species as well (Tea et al., 2012).

Egg cold hardiness has been much less studied for the Asian populations: it is generally accepted that the Asian GM populations are more cold-resistant than the European ones (Pantyukhov, 1964). This point of view seems plausible, for the Asian populations in Russia are found in the areas with harsh continental climate. According to long-term meteorological observations (www.pogodaiklimat.ru), in the range of the Siberian GM populations, air temperature repeatedly drops to –30…–42 °C, whereas monthly mean temperatures are –10.~ 30 °C. However, our results show that the lower cold hardiness limit of the Asian populations (– 29.0…–29.9 °C) lies very close (to within one degree) to that of the European populations. This agreement indicates that the lower lethal temperature for *L. dispar* eggs is conservative. The obtained results promote a new vision of factors which determine the northern range of gypsy moth in Asia. It turns out that the West Siberian and the East Siberian geographical forms of GM are almost exclusively found in territories where the temperature repeatedly falls below –30 °C (and sometimes below –40 °C) during the winter, thereby going beyond the physiological tolerance limits of eggs. Thus, extreme winter air temperatures themselves cannot be a factor constraining the northbound *L. dispar* asiatica range expansion. Ovipositional behavior is most variable over the wide distribution of this subspecies: members of each geographical form prefer different biotopes (Pogue and Schaefer 2007). For example, GM females of the West Siberian geographical form, including those from the Novosibirsk population, will preferably oviposit on trees at the average height of 6-7 cm above the ground (Ponomarev et al., 2012). Ovipositing so close to the ground, females ensure that they will be covered with snow. This behaviour is entirely rational, because the temperature under the snow is substantially higher are more stable compared to the air temperature (Leonard 1974; Madrid and Stewart, 1981). The role of snow cover in the winter survival of GM eggs is pointed out in Nealis et al. (2012). Pogue and Schaefer (2007) considered different arrangements of GM eggs, some of which are probably designed to better cover the egg masses under the snow. Thus, the presence and stability of snow cover is a critical factor favouring the survival of the eggs of the West Siberian geographical form. On the contrary, there are no restrictions of this kind in the behaviour of females of the Central Asian geographical form (Jalal-Abad region of the Kyrgyz Republic): their eggs are found on stones and trees at heights up to several metres (Romanenko, 1981; Ponomarev et al., 2012; the authors’ own observations). Such non-selectivity is probably due to the fact that according to long-term meteorological observations (www.pogodaiklimat.ru), the temperature has never fallen below –27 °C in the range of the Kyrgyz population, and the average winter temperatures are around 0 °C.

Apart from the behavioural adaptation, we have found that the Siberian populations are different from the Central Asian one in that they are more resistant to prolonged exposure to the sublethal temperature (–20 °C). We believe that these forms of physiological adaptation can be useful in late autumn: in November, the temperature in Siberia can fluctuate from above-freezing values to –30 °C, with no snow cover. The lesser cold hardiness of the eggs of the Kyrgyz population during the prolonged exposure at –20 °C can apparently be explained by that it inhabits areas with milder climate. Thus, the decisive factor that enabled GM to colonize vast territories of West and East Siberia is apparently the high flexibility of ovipositional behavior of gypsy moth females combined with a number of physiological adaptations.

The importance of different factors determining patterns of GM population outbreaks is debatable. No single hypothesis can explain all peculiarities of forest Lepidoptera population outbreak dynamics, and only the combination of several factors can provide an acceptable explanation (Myers and Cory, 2013; Elderd et al., 2013; Páez et al., 2015). The results of the present study support the hypothesis of possible direct influence of weather on outbreak patterns. Indeed, the temperature in the range of the Siberian GM populations repeatedly drops below – 30 °C in winter, this is why GM eggs can only survive under the snow cover. Hence, it can be supposed that the insect outbreak dynamics can be affected by the date of the formation of snow cover. For example, if at the beginning of winter there is a serious cold snap before the egg masses are covered with snow, then most GM eggs will die, which can contribute to the end of the GM population outbreak. On the contrary, an early formation of snow cover (before severe frosts), along with other factors, can provoke a GM population outbreak, because most eggs will successfully overwinter. We believe that one of the causes of the observed asynchronicity of GM population outbreaks in the Asian part of its range (Martemyanov et al., 2019) may be fluctuations of weather conditions. It is in contrast to the European part of the GM range, where the outbreaks were synchronous (Liebhold et al., 2000; Esper et al. 2007; Soukhovolsky et al., 2015) and where winter temperatures are usually not below –30 °C.

Based on available data on the egg cold hardiness of the European populations of *L. dispar dispar* and the results of the present study, we can assume that the physiological tolerance limit of the *L. dispar* eggs is not below –30…–31 °C. This conclusion is not in agreement with results by Pantyukhov (1964), obtained during the experiments of 1958–1960. Low-temperature freezers were then unavailable, this is why the author used a mixture of solid carbon dioxide (dry ice) with water to obtain temperatures of –45…–48 °C. It is an unstable heterogeneous system, in which it is difficult to maintain a given temperature. These technical issues could lead to overestimated cold hardiness of the North Caucasian and Far Eastern GM populations, which were studied by Pantyukhov. He states that “North Caucasian gypsy moth eggs die almost completely after 12–14 hours of cooling, whereas the Far Eastern eggs — after 22–24 hours”. Unfortunately, we were unable to test eggs of the Far Eastern GM population for cold hardiness. But it is known that extreme and moderate winter temperatures on the Far East are close to those in Siberia. Notably, egg masses of the Far Eastern GM population winter in the forest floor. It has been observed (Yurchenko and Turova, 1984) that eggs wintering in the forest floor, are characterized by high viability (up to 80–90 %), whereas eggs laid on fences and poles often die in winter. It can be therefore supposed that the Far Eastern GM geographical form is not more cold-resistant than the Siberian population. It also follows from the above citation that the eggs of the North Caucasian population die earlier than those of the Far Eastern population, but still a small proportion of them survives up to 12 hours of exposure at –45…–48 °C. This statement sounds doubtful, because the North Caucasian region is among the warmest in Russia: only two times over the last 50 years of meteorological observations the local temperature fell below – 30 °C, and it never happened in most North Caucasian regions. This is why the egg cold hardiness reported by Pantyukhov appears superfluous for the North Caucasian population.

In general, almost the entire range of *L. dispar asiatica* in Russia falls on territories where the winter temperatures go beyond the limits of egg physiological stability. In the conditions of the harsh continental climate of the Northern Asia, the lack of physiological tolerance is compensated by the flexibility of ovipositional behavior of females, and *L. dispar asiatica* survives thanks to the use of warmer (compared to the air temperature) biotopes for oviposition. The case of the West Siberian geographical form shows that in presence of stable snow cover, extreme winter temperatures cannot constrain the northbound expansion of the GM range. Forecasts of GM range changes should take into account not only the winter temperature, but also dates of formation, stability and depth of snow cover (Sullivan and Wallace, 1972; Smitley et al., 1998; Streifel et al., 2019).

To conclude, let us make several remarks regarding the techniques for the assessment of insect cold hardiness. Our experience in field research suggests that the conclusions concerning the temperature stability of GM eggs must be based on the reads of local temperature loggers, whose detectors are in close contact with egg masses. This is the only way to ensure the completeness and accuracy of data on the effects of extreme temperatures on egg samples. Failure of this condition can result in serious errors. Data from local weather stations can be used only when average (full-day, full-month) temperatures are in question. But when it comes to the extreme temperature of samples located in a particular biotope, the data of weather stations can only serve as a guide. For example, during the field experiment No. 1 (December 28, 2019) we recorded the minimum temperature of –30.2 °C; at the same moment the minimum reads from the three nearest weather stations (within a radius of 20 km) were –29.0 °C, –31.6 °C, and – 34.5 °C, respectively. Besides the air temperature, extreme local temperatures depend on a number of additional factors, such as height above the ground, site illumination, the presence of snow cover etc. The presence of snow has a particularly strong effect: the difference in egg temperatures above and below the level of snow cover can measure up tens of degrees (see Fig. 2). Therefore, when making conclusions about the viability of eggs after wintering, it is necessary to make sure whether the egg masses wintered under the snow or above the level of snow cover. The data recording frequency is also important: the recording interval should preferably not exceed 1-2 hours, because with longer intervals, the extremum point of daily temperature curve can be missed.

## List of symbols and abbreviations

GM: Gypsy moth

## Author contributions

A.G.G and K.A.V. contributed equally to this work.

